# Single-cell 5’ RNA sequencing of camelid peripheral B cells reveals cellular basis of heavy-chain antibody production

**DOI:** 10.1101/2023.11.30.569338

**Authors:** Li Yi, Xin Guo, Yuexing Liu, Jirimutu, Zhen Wang

**Affiliations:** Key Laboratory of Dairy Biotechnology and Engineering, Ministry of Education; College of Food Science and Engineering, Inner Mongolia Agricultural University, Huhhot, China; CAS Key Laboratory of Computational Biology, Shanghai Institute of Nutrition and Health, University of Chinese Academy of Sciences, Chinese Academy of Sciences, Shanghai, China; Guangzhou Laboratory, Guangzhou, China; Inner Mongolia China-Kazakhstan Camel Research Institute, Alxa 750306, China

**Author notes:** Correspondence (J.), (Z. Wang). These authors contributed equally (Li Yi, Xin Guo).

## Abstract

Camelids are capable of producing both conventional tetrameric antibodies (Abs) and dimeric heavy-chain antibodies (HCAbs). While B cells generating these two types of Abs exhibit distinct B-cell receptors (BCRs), it remains unclear whether these two B cell populations differ in their phenotypes and developmental processes. Here, we collected eight PBMC samples before and after immunization from four Bactrian camels and conducted single-cell 5’ RNA sequencing. We characterized the functional subtypes and differentiation trajectories of circulating B cells in camels, including native B cells, memory B cells, intermediate B cells, atypical B cells, and plasma cells. Additionally, we reconstructed single-cell BCR sequences and revealed the IGHV and IGHC gene types. We found that B cells expressing variable genes of HACbs (VHH) were widely present in various functional subtypes and showed highly overlapping differentiation trajectories to B cells expressing variable genes of conventional Abs (VH). After immunization, the transcriptional changes in VHH+ and VH+ B cells were also largely consistent. Our study elucidates the cellular context of HCAb production in camels, and lays the foundation for the development of single B cell-based nanobody screening.

## Introduction

It is well recognized that conventional antibodies (Abs) or immunoglobulins (Igs) in vertebrates are heterotetramers composed of two heavy chains and two light chains [1]. In camelids, however, there are also homodimeric IgGs containing only two heavy chains, known as heavy chain antibodies (HCAbs) [2]. Despite the absence of light chains, the HCAb has unique mechanisms for generating structural diversity. Compared with conventional Abs, HCAbs may have a longer third complementarity determining region (CDR3) and may have more cysteine residues within the variable regions to form additional disulfide bonds [3]. The single-domain Ab engineered by retaining only the variable region of the HCAb is called a nanobody because of its extremely small molecular weight [3]. Nanobodies not only maintain antigen binding affinity, but also have higher stability, hydrophilicity and tissue permeability, so they have been more and more widely used in biomedicine [4]. In 2018, Caplacizumab, the first nanobody drug developed by Ablynx, has been approved in Europe [5].

Genetically, HCAbs are encoded by dedicated genes that differ from those encoding conventional Abs in two ways. One is that in the constant region gene, IGHG2 and IGHG3 encoding the HCAb have splice site mutations (GT->AT) at the exon of the first constant domain of the heavy chain (CH1), resulting in the deletion of CH1 connected to the light chain [6–8]. Neither IGHG1 nor other IGHC genes (including IGHM, IGHE, IGHA) encoding conventional Abs had this mutation [8]. The second is that the IGHV gene of the HCAb undergoes characteristic amino acid substitutions at four positions in the second framework region (FR2), most of which are: V42F, G49E, L50R and W52G [9–11]. In conventional Abs, these four positions are involved in the binding of the light chain, so they are highly conserved hydrophobic amino acids (called VH). In HCAbs, they are usually replaced by hydrophilic amino acids (called VHH). Sequencing and assembling of the camelid IGH locus showed that both HCAbs and conventional Abs are encoded by genes in the same IGH locus with a typical V-D-J-C organization, and their coding genes are intermixed within the IGHV and IGHC regions [12, 13]. The IGHD and IGHJ genes are common to both conventional Abs and HCAbs [12, 13]. Immune repertoire sequencing of multiple camelids reveals genetic diversity of VHH, both at the germline and rearranged level [11, 12, 14, 15].

Studies on model animals have shown that the differentiation process of B cells has undergone multiple stages of change [16, 17], and the functional subsets of these B cells are mainly distinguished by the rearrangement status of B cell receptor (BCR) gene segments and cell surface markers [18]. The co-existence of B cells producing conventional Abs and HCAbs in camelids raises the question of whether these two populations of B cells have gone through similar developmental stages. Although B cells generating HCAbs were speculated to pass through a naive IgM+ stage similar to the case for conventional Abs [12], the cellular components remain largely unexplored due to the lack of surface markers and corresponding reagents for camelids.

In recent years, the development of single-cell RNA sequencing (scRNA-seq) technology has led to a revolution in the study of cell heterogeneity [19, 20]. One of the greatest advantages of scRNA-seq is the ability to define cell types at the genomic level, allowing unbiased phenotyping of all cells. This is particularly useful for non-model organisms where traditional experimental tools are limited [21–23]. A study performed scRNA-seq for peripheral blood mononuclear cells (PBMCs) of an immunized alpaca and described alterations in cell type composition and gene expression [24]. However, BCR types cannot be recovered with 3’ RNA sequencing data, making it impossible to distinguish B cells expressing conventional Abs and HCAbs.

In this study, we performed single-cell profiling of PBMCs of four Bactrian camels using 5’ RNA sequencing. First, we identified cell types in the camel PBMCs, especially B cell subtypes and differentiation trajectories. We then reconstructed the BCR sequences for each B cell, distinguishing IGHV and IGHC types and revealed their correspondence with B cell phenotypes. Finally, we compared changes in PBMCs and B cells between pre- and post-immunization. Our results showed that while B cells producing conventional Abs and HCAbs displayed preference in BCR sequences, their differentiation trajectories largely overlapped. The transcriptional changes of the two B cell populations were also highly similar through the immunization process.

## Results

### Cell composition of camel PBMCs

We isolated PBMCs from four healthy adult Bactrian camels (C1-C4, Supplementary Table 1), and performed scRNA-seq for the PBMCs based on the 10× Genomics platform. In order to recover IGHV and IGHC types of BCRs, 5’ RNA libraries were constructed for sequencing. About 5,500-10,000 cells were detected per sample, and the median genes detected per cell were 1,200-1,700 (Supplementary Table 2). After quality control by UMI counts and mitochondrial percentages, a total of 26,759 cells were preserved.

We integrated cells across all samples with Seurat [25], and performed dimensional reduction and unsupervised clustering (Fig. 1A). Major cell types of camel PBMCs could be recognized with the expression of canonical marker genes as in humans (Fig. 1B), including CD4+ T cells (CD3, CD4, 24.24%), CD8+ T cells (CD3, CD8, 14.41%), γδ T cells (CD3, TRDC, 7.67%), profiling T cells (CD3, TYMS, 2.51%), natural killer (NK) cells (KLRB1, NKG7, FCGR3A, 1.84%), B cells (CD19, MS4A1, 33.59%), plasma cells (CD38, TYMS, 2.67%), monocytes (CD14, CD68, 11.96%) and dendritic cells (ITGAX, FCER1A, IL3RA, NRP1, 1.12%). The four camels showed similar cell compositions (Supplementary Fig. 1). The lymphocyte composition of CD4+ T cells, γδ T cells and B cells was comparable to previous studies based on flow cytometry [26], although other lymphocytes could not be identified by flow cytometry due to the lacking of cross-reactive monoclonal antibodies. Notably, camels belong to γδ-high mammalian species as other artiodactyls [26], in contrast to γδ-low species like humans and mice (<5% in PBMCs [27]).

**Figure 1.**
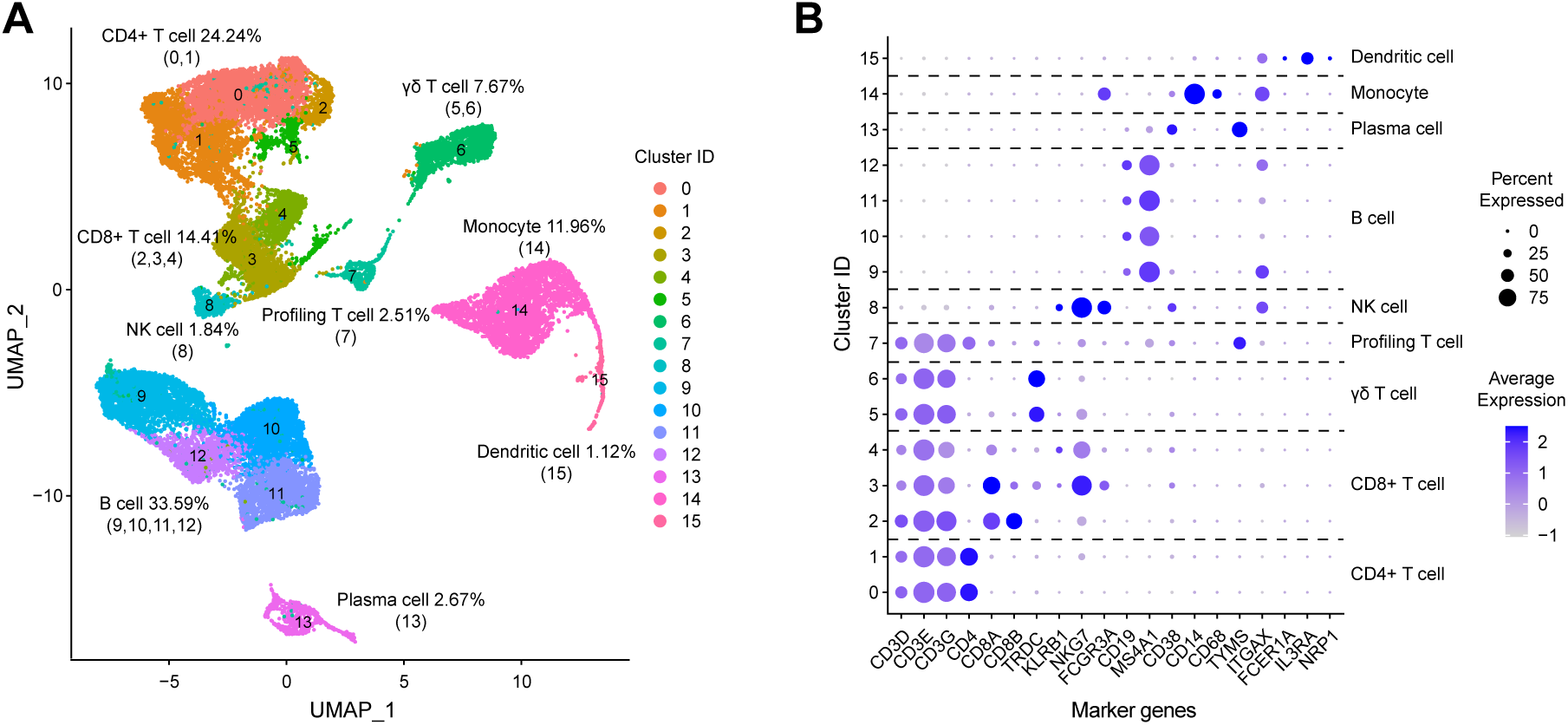
Major cell types of camel PBMCs revealed by scRNA-seq. (A) UMAP visualization of cell clustering. Cell types are annotated based on the expression of marker genes and their proportion in PBMCs is calculated. (B) Dot plot depicting average expression and expression percentage of marker genes in each cell cluster.

Our scRNA-seq analysis also enabled to dissect the heterogeneity of major cell types with high resolution. For example, Cluster 0 (CD4+ T), 2 (CD8+ T), and 5 (γδ T) showed high expression of CCR7 and SELL but low expression of S100A4, suggesting a phenotype of naive T cells [28] (Supplementary Fig. 2). Cluster 1 (CD4+ T) and 6 (γδ T) showed high expression of S100A4, and Cluster 3/4 (CD8+ T) showed high expression of GZMK, suggesting a phenotype of memory T cells [28] (Supplementary Fig. 2).

### B cell subsets and differentiation trajectories

As Abs or BCRs were expressed by B cells, we performed an in-depth analysis of B cell subtypes. Apart from plasma cells (Cluster 13, 7.35%), unsupervised clustering divided B cells into four subtypes (Cluster 9-12, Fig. 2A), which showed similar proportion in the four camels (Supplementary Fig. 3). We first inferred their functional annotations based on differentially expressed genes among the subtypes (Fig. 2B). Cluster 9 (31.58%) had a phenotype of atypical B cells [29, 30], with a relatively higher expression of ITGAX and TBX21, but a relatively lower expression of CR2. Unlike in humans where only a small amount of peripheral atypical B cells was present, camels had a high proportion of atypical B cells, which was observed in other species such as horses [22]. Cluster 10 (26.43%) expressed higher levels of CXCR4, CD69 and SELL, which was similar to human naive B cells [30, 31]. Cluster 11 (20.42%) expressed higher levels of ITGB1, ITGB7, S100A6, S100A13 and IGHA, which were related to functions of memory B cells [30]. Cluster 12 (14.22%) showed a higher expression of IGHM. We also mapped the camel B cell transcriptome to the human PBMC reference dataset built in Seurat [25]. The result indicated that, except for the lacking of atypical B cells in the human dataset, Cluster 10, 11 and 12 had the highest similarity to human naive, memory and intermediate B cells, respectively (Supplementary Fig. 4). Therefore, we assigned corresponding cell type labels to these clusters (Fig. 2A). Notably, CD27, a marker gene for human memory B cells [18], was expressed at low levels in all camel B cell subsets (Supplementary Fig. 5). IGHD, a marker gene for human naive B cells [18], also lacked expression and it was previously identified as a pseudogene in camelid genomes [12, 13].

**Figure 2.**
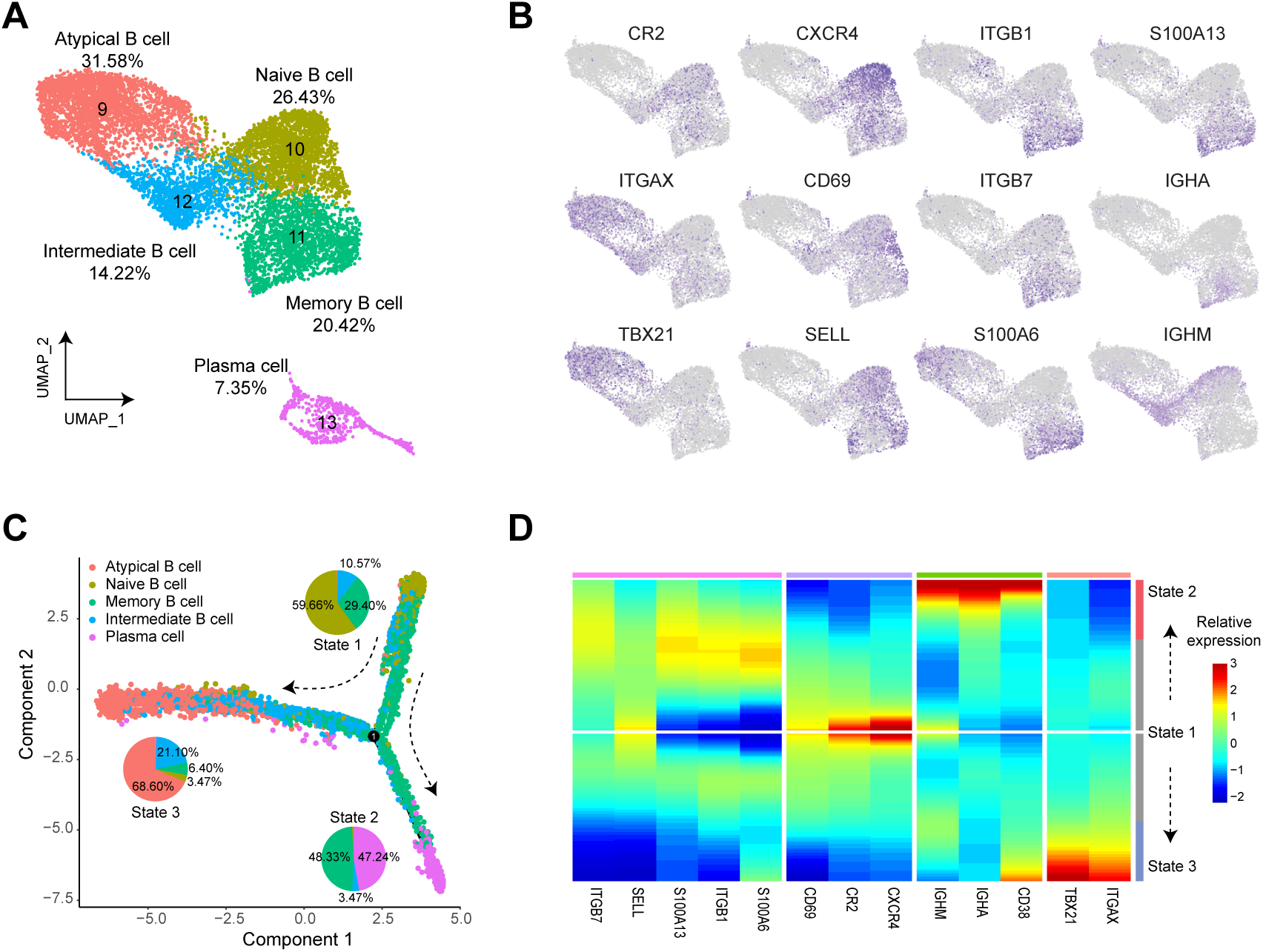
Characterization of camel B cell subtypes. (A) UMAP plot of B cells colored by their subtypes. The proportion of each subtype in B cells is calculated. (B) Expression of marker genes among the B cell subtypes identified by Seurat FindMarkers. Atypical B cells show relatively lower expression of CR2 but higher expression of ITGAX and TBX21. Naive B cells show higher expression of CXCR4, CD69 and SELL. Memory B cells show higher expression of ITGB1, ITGB7, S100A6, S100A13 and IGHA. Intermediate B cells show higher expression of IGHM. (C) B cell state and differentiation trajectories inferred with Monocle2. Cells are colored according to their subtypes, and the proportion of subtypes in each state is displayed by pie plot. The pseudotime is assigned by assuming naive B cells as the root. (D) Heatmap depicting the relative expression of marker genes along the trajectories.

To explore the relationships between these B cell subtypes, we used Monocle2 [32] to infer the single-cell trajectories. Monocle2 identified three B cell states (Fig. 2C): State 1 consisted mostly of naive B cells (59.66%) and memory B cells (29.40%); State 2 consisted mostly of memory B cells (48.33%) and plasma cells (47.24%); State 3 consisted mostly of atypical B cells (68.60%) and intermediate B cells (21.10%). Moreover, starting from State 1, Monocle2 revealed two major differentiation trajectories of B cells: 1) naive B cells - memory B cells - plasma cells; 2) naive B cells - intermediate B cells - atypical B cells. Similar to humans [33], this result suggested that atypical B cell differentiation was an alternative lineage distinct from classical lineage in camels. The expression trends of marker genes for the B cell subtypes were consistent with these two differentiation trajectories (Fig. 2D).

### BCR types and association with B cell subsets

We first determined the IGHC type of BCRs based on the single-cell expression matrix (Fig. 3A and Supplementary Fig. 6). Among all B cells, 31.63% had their expressed IGHC type recovered, with the highest recovery rate in plasma cells (89.76%) due to the fact that plasma cells had the highest BCR expression levels. Consistent with the developmental stages of B cells, both naive and intermediate B cells primarily expressed IGHM without undergoing class switching, while memory B cells, atypical B cells and plasma cells mainly expressed IGHG and IGHA that had undergone class switching (Fig. 3A). The plasma cells expressing HCAb genes (IGHG2/3+) accounted for 60.79% of all IGHG+ plasma cells, in agreement with the fact that HCAbs were the major IgG form in the serums of Bactrian camels [34]. We also identified the type of light chain genes expressed in each B cell and found that even for IGHG2/3+ B cells, there was obvious expression of IGLC or IGKC (Supplementary Fig. 7). This indicated that the absence of light chains in HCAbs was not due to suppression of light chain gene transcription.

**Figure 3.**
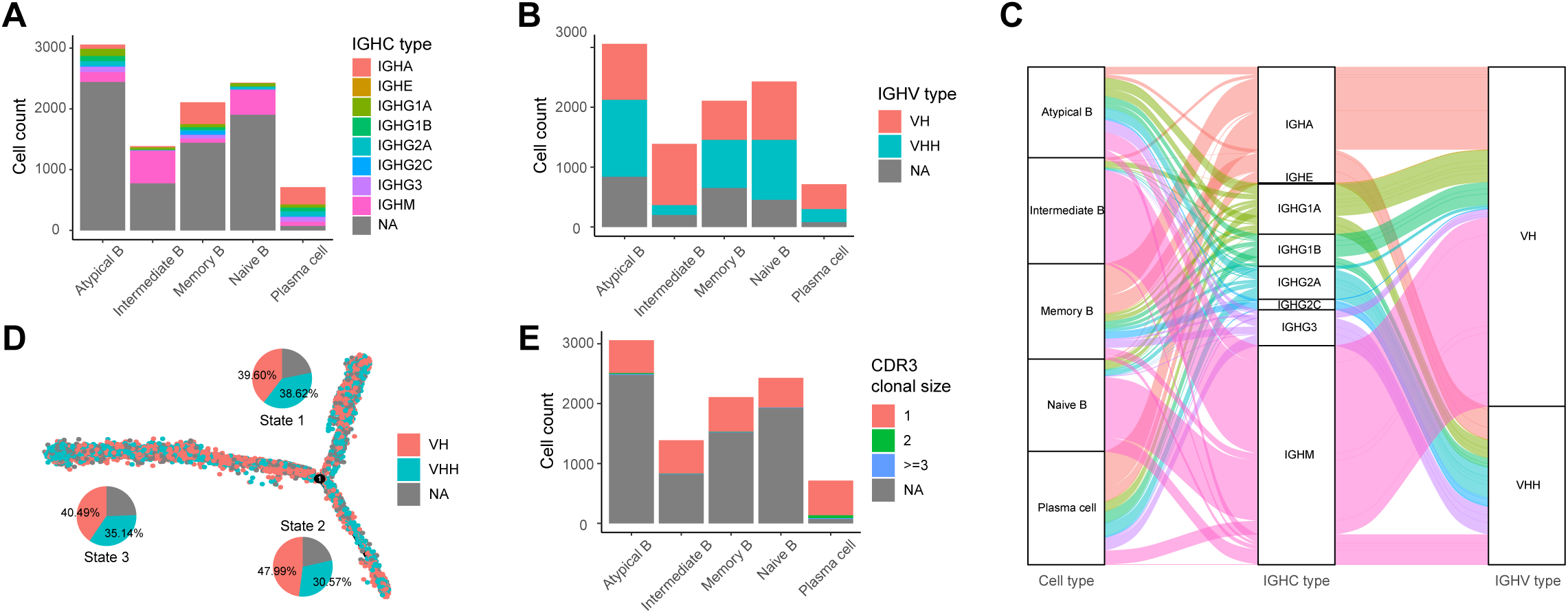
Characterization of camel single-cell BCRs. (A) Number of cells with different IGHC types in each B cell subtype. The IGHC types are assigned based on the gene expression. NA represents cells without IGHC expression detected. (B) Number of cells with different IGHV types (VH/VHH) in each B cell subtype. The IGHV types are assigned based on the hallmark substitutions in FR2. NA represents cells without FR2 recovered. (C) Sankey diagram depicting the relationship between B cell subtypes, IGHC types and IGHV types. Although most IGHG2/3+ cells are associated with VHH, and most conventional IGHC are associated with VH, non-typical associations are widely present. (D) Mapping IGHV types to differentiation trajectories of B cells inferred with Monocle2. Cells are colored according to their IGHV types, and the proportion in each state is displayed by pie plot. (E) Number of cells with different clonal size in each B cell subtype. The clonotypes are determined by IGHV CDR3 sequences. NA represents cells without CDR3 recovered.

To determine the variable region sequence of the BCRs, we used TRUST4 [35] to assemble the reads mapped to the IGH locus at the single-cell level. The reference IGH sequences were obtained from our previously published Bactrian camel genome [13]. For contigs assembled, further annotation was performed based on IgBLAST [36], including IGHV/D/J/C genes, FR and CDR positions and sequences. The IGHC types determined by sequence assembly and UMI count were highly consistent (Supplementary Fig. 8). We divided B cells into VH+ and VHH+ types based on the characteristic amino acid substitutions in the FR2 region of IGHV genes (Supplementary Fig. 9). 77.09% of all B cells had their IGHV type recovered (Supplementary Fig.10), and each B cell subset contained a comparable proportion of VH+ and VHH+ cells, with only a lower proportion of VHH in intermediate B cells (Fig. 3B). Furthermore, we generated a Sankey diagram showing the relationship between B cell subsets, IGHC types and IGHV types (Fig. 3C). As expected, the majority of IGHG2/3+ B cells were paired with VHH, though a small proportion of IGHG2/3+VH+ B cells was also identified. This was consistent with the fact that some HCAbs were isolated with an antigen-binding VH domain [3, 37]. Similarly, most IGHG1+, IGHM+, and IGHA+ B cells were paired with VH, though VHH could also contribute to the conventional BCRs (Fig. 3C). These non-typical BCRs could be expressed by plasma cells, indicating that they were not only a transitional state during class switching but might also have specific antigen binding capabilities. We also mapped the IGHV type to the differentiation trajectories of B cells reconstructed by Monocle2 (Fig. 3D). The VHH+ and VH+ cells were well mixed in terms of differentiation states and trajectories, indicating that they underwent similar developmental process.

We analyzed the diversity of reconstructed heavy chain CDR3 sequences. Among all B cells, 29.87% had complete CDR3 sequences recovered, while this proportion reached 90.18% in plasma cells (Fig. 3E and Supplementary Fig. 11). As expected, compared to VH, the CDR3 region of VHH was significantly longer [3] (Supplementary Fig. 12), and the sequence features were similar to previous reports [38, 39] (Supplementary Fig. 13). The vast majority of B cells (95.31%) had unique CDR3 sequences (i.e., clone size = 1), and the unique proportion only slightly decreased in plasma cells (90.05%), indicating that naturally occurring CDR3 sequences were highly diverse in camels (Fig. 3E).

### Comparative analysis before and after immunization

Camel immunization was an important way to prepare antigen-specific nanobodies. To elucidate the changes in PBMCs and B cells following immunization, we immunized two of the Bactrian camels (C3, C4) with bovine serum albumin (BSA). Immunization was performed every two weeks, and enzyme-linked immunosorbent assay (ELISA) indicated that the antibody titer reached the highest level after four rounds of immunization (Fig. 4A). We performed scRNA-seq on PBMCs collected at 42 and 56 days post-immunization (Supplementary Table 3) and compared them with the pre-immunization samples (0 day). We transferred cell-type annotations from our previously analyzed samples to the post-immunization samples (Supplementary Fig. 14). The proportions of most cell types showed no significant changes (Supplementary Fig. 15), except for a significant increase in the proportion of γδ T cells after immunization (*P* = 0.024, two-sided *t*-test).

**Figure 4.**
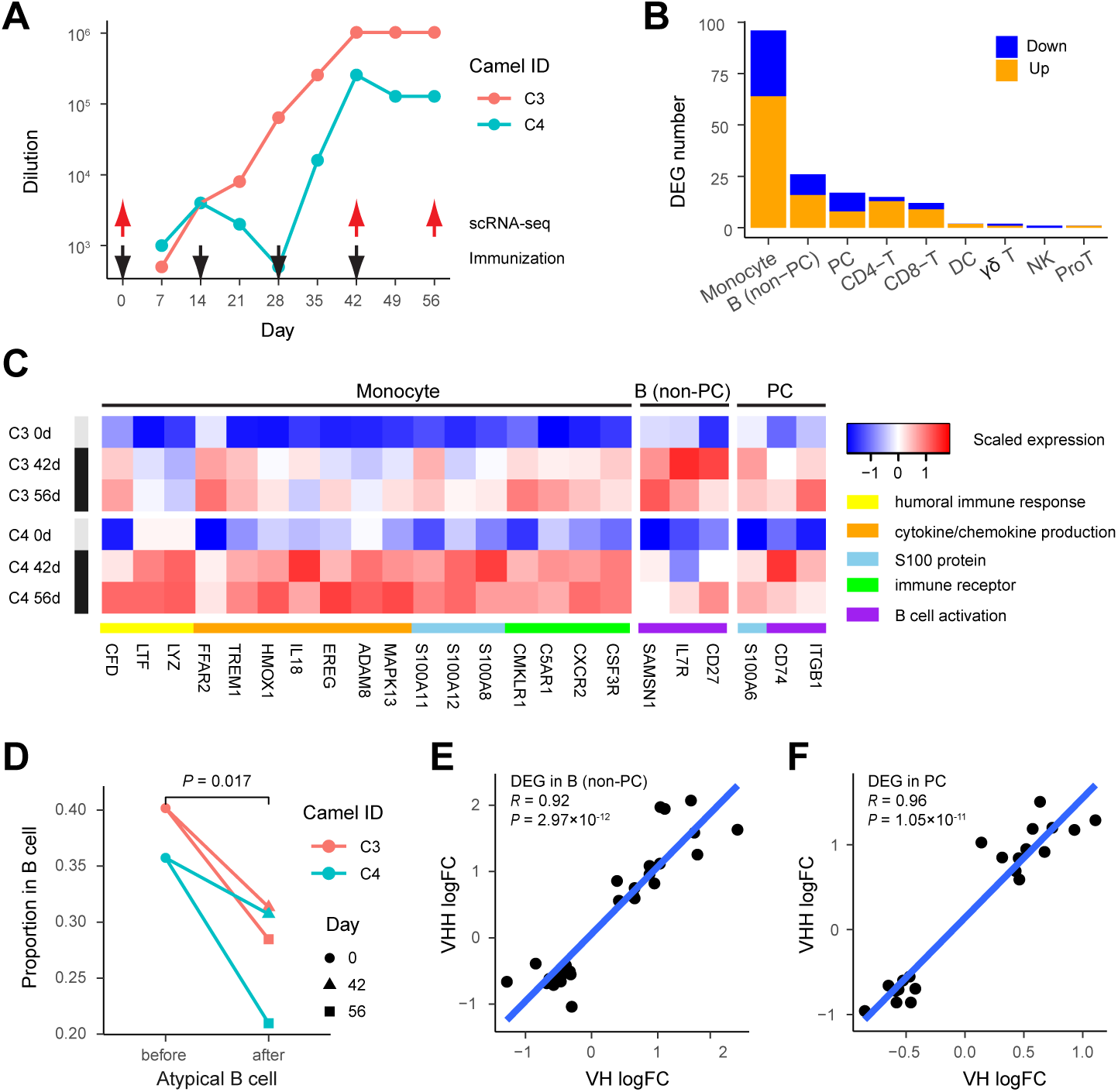
scRNA-seq of post-immunization PBMCs and comparative analysis. (A) Immunization schedule and serum antibody titer. Two camels received four doses of BSA on the days indicated by black arrows, and PBMCs were collected for scRNA-seq before and after immunization on the days indicated by red arrows. The antibody titer was measured as the highest serum dilution using ELISA. (B) Number of DEGs identified in major cell types. Up-regulated genes in post-immunization samples compared to pre-immunization samples are colored in orange, while down-regulated genes are colored in blue. (C) Heat map of selected DEGs with functional enrichment in monocytes and B cells (PC, plasma cell). Gene expression on the pseudobulk level is normalized by the counts per million mapped reads and scaled across samples. (D) Proportion of atypical B cells in all B cells before and after immunization. *P*-values are calculated using the two-sided *t*-test. (E, F) Comparison of the log fold change (logFC) of DEGs between VH+ and VHH+ B cells during immunization. DEGs are identified in (E) non-PCs and (F) PCs, respectively. Pearson’s correlation coefficient *R* and its *P*-value are calculated.

For each major cell type, we conducted differential gene expression analysis between pre- and post-immunization using the pseudobulk approach [40]. Due to the higher average gene expression levels in plasma cells, B cells were divided into non-plasma cells and plasma cells for pseudobulk analysis, respectively. We identified the highest number of differentially expressed genes (DEGs, adjusted *P* < 0.05) in monocytes, followed by B cells and plasma cells, with an overall upregulation of DEGs after immunization (Fig. 4B). Furthermore, we performed functional enrichment analysis on the DEGs. The upregulated genes in monocytes were significantly enriched in biological processes and functions related to humoral immune response, cytokine/chemokine production, S100 proteins, and immune receptors, consistent with the activation of inflammatory responses during immunization (Fig. 4C). The significantly upregulated genes in B cells and plasma cells were enriched in B cell activation process, consistent with the increased serum antibody titer.

Comparison of B cell subtypes showed a significant decrease in the proportion of atypical B cells after immunization (*P* = 0.017, two-sided *t*-test, Fig. 4D and Supplementary Fig. 16), suggesting that immunization may reduce the differentiation of B cells into atypical B cells. Accordingly, there was a slight decrease in the proportion of VHH+ B cells after immunization (*P* = 0.040, two-sided *t*-test, Supplementary Fig. 17). However, our BCR analysis of the post-immunization samples did not reveal significant changes in IGHC proportions (Supplementary Fig. 18). We also did not observe significant changes in clonotype diversity (Supplementary Fig. 19), possibly due to different kinetics in plasma cell abundance and serum antibody concentration [24, 41]. To determine whether VH+ and VHH+ B cells exhibit consistent transcriptional responses before and after immunization, we conducted differential expression analyses using the pseudobulk method for VH+ and VHH+ B cells, respectively. The log fold changes of DEGs between VH+ and VHH+ B cells exhibited a strong positive correlation, which was observed in both non-plasma cells (Fig. 4E) and plasma cells (Fig. 4F). These results indicated a highly similar transcriptional response of VH+ and VHH+ B cells during the immune process.

## Discussion

Camelid species are the only known mammals capable of producing HCAbs. The sequence features and genetic basis of camelid HCAbs have been well characterized and widely applied in nanobody engineering [3, 4]. However, the cellular process underlying the production of HCAbs remains poorly understood. It was observed that certain IgM+ B cells could express VHH genes without somatic mutations, suggesting that B cells bearing dimeric IgG might undergo a naive IgM+ stage, similar to the case for conventional IgG-producing cells [12]. However, the phenotypes of camelid B cells were not fully characterized due to the absence of surface markers and corresponding reagents [26]. Recently, scRNA-seq was performed on PBMCs of an alpaca during immunization process [24], but 3’ RNA sequencing was unable to distinguish the BCR types. In this study, we employed single-cell 5’ RNA sequencing to simultaneously reveal the phenotypes and BCR sequences of peripheral B cells in Bactrian camels.

First, we identified cell components in camelid PBMCs, especially B cell subtypes. Compared with humans and mice, camel PBMCs contained a higher proportion of γδ T cells and atypical B cells, providing a unique source for studying the functions of these cells. Our pseudotime analysis revealed two differentiation trajectories of the B cells. In addition to the classical trajectory involving naive B cells, memory B cells and plasma cells, there was also an alternative trajectory involving naive B cells, intermediate B cells and atypical B cells. Interestingly, it was recently reported that atypical B cells were part of an alternative lineage of B cells in humans, which could participate in responses to vaccination and infection [33].

By reconstructing the BCR sequences, we could determine the IGHV (VHH+, VH+) and IGHC gene types (IGHG2/3+, IGHG1, and others) of each B cell. We observed the presence of VHH+ B cells in all functional subtypes, and their differentiation trajectories largely overlapped with VH+ B cells, indicating that both VHH+ and VH+ cells experienced similar developmental stages. While VHH genes were known to preferentially pair with IGHG2/3 to form HCAbs, and VH genes typically pair with other IGHC genes to form conventional Abs, we found widespread presence of BCRs with non-typical connections between IGHV and IGHC genes. These non-typical connections may have two implications. One was a transitional state of IGHC class switching. Indeed, in naive B cells, VHH-bearing cells could pair with IGHM, resulting in an IgM+ state [12]. Additionally, these BCRs with non-typical connections could possess specific functions. In fact, in nanobody cloning, HCAbs with VH genes and high antigen affinity were isolated [37]. We also found that camel plasma cells could express BCRs in the form of VHH+IGHA+. The protein structure and function of these BCRs were worthy of further investigation.

We also investigated the dynamic changes in camel PBMCs before and after BSA immunization. We found that monocytes exhibited the most significant transcriptional changes post-immunization, characterized by upregulation of inflammation-related genes. B cells also showed activation signals, and the transcriptional changes in VHH+ and VH+ cells were highly consistent. This indicated that the transcriptional regulatory programs of the two B cell populations did not exhibit significant differences. It was worth noting that we sequenced cells at the time of the highest serum antibody titer, which may not necessarily correspond to the highest plasma cell abundance [41]. In fact, Lyu et al. [24] found that the proportion of plasma cells in an immunized alpaca reached its peak 3 days after the second antigen simulation, but rapidly returned to the pre-immunization level 2 days later. Moreover, in antibody discovery, antigen-specific B cell sorting combined with single-cell BCR sequencing is a rapid screening strategy that has been successfully applied in humans [42] and animals [43]. The camel BCR reconstruction method we have developed can also be used in this scenario to facilitate rapid nanobody discovery.

## Methods

### Camel PBMC collection

We chose two male and two female healthy Bactrian camels aged 4-5 years from Tuzuo Banner, Inner Mongolia, China. For each camel, 50 ml peripheral blood was collected from the jugular vein and placed in sodium citrate anticoagulation tubes after disinfection treatment. The collections were made following the guidelines from the Camel Protection Association of Inner Mongolia. PBMCs were isolated from the peripheral blood with density gradient centrifugation using the Ficoll-Paque medium. 3ml PBMCs were transferred into 5ml cryotubes with 2ml cell freezing medium (Yeasen, 40128ES50). Cells were stored frozen at -80°C for one day and then transported to sequencing lab on dry ice.

### Single-cell library construction and sequencing

The viability of thawed cells was examined to exceed 80% with trypan blue staining. Single-cell libraries were constructed using the Chromium Next GEM Single Cell V(D)J Reagent Kits v1.1 (10x Genomics, 5’ Library Kit) according to the manufacturer’s instructions. Specifically, cells, barcoded gel beads and partitioning oil were combined on a microfluidic chip to form Gel Beads-in-Emulsion (GEMs). Within each GEM, a single cell was lysed and the transcripts were identically barcoded through reverse transcription. The cDNA libraries were sequenced on an Illumina NovaSeq platform to generate 2×150-bp paired-end reads.

### scRNA-seq data analysis

Sequencing reads were processed by the Cell Ranger pipeline (v6.1.2, 10x Genomics). Genome alignment was performed against the assembly BCGSAC_Cfer_1.0 (GCF_009834535.1). Only annotations of protein-coding genes in RefSeq, including the mitochondrial genes were preserved. Our previous BCR/TCR gene annotations [13] were also incorporated as they were incomplete in RefSeq. The gene-barcode count matrix of unique molecular identifier (UMI) was analyzed with Seurat (v4.0.0) [25]. Specifically, cells with total UMI count between 1,000 and 60,000, and mitochondrial gene percentage <5% were preserved for quality control. The count matrix was normalized using sctransform [44], and cells across different samples were integrated with the canonical correlation analysis (CCA). The principal component analysis (PCA) was applied for the integrated matrix, and the top 30 PCs were then used for the uniform manifold approximation and projection (UMAP) and the nearest neighbor graph-based clustering. The resolution of clustering was set at 0.5. Marker genes were identified with FindMarkers between clusters. We also mapped the camel B cells to the human reference PBMC dataset using Seurat’s reference mapping utility [25].

### Differentiation trajectory inference

The differentiation trajectories among B cells were inferred with Monocle2 (v2.18.0) [32]. The UMI count matrix was modeled with the negative binomial distribution, and size factors and dispersions were estimated. Genes expressed in at least 10 cells were preserved. We chose genes with high dispersion across cells to define the trajectory, which had mean expression > 0.1 and dispersion greater than the fitted value. DDRTree was performed for dimensional reduction, and cells were ordered in the reduced dimensional space. We assigned pseudotime to the trajectories by assuming naive B cells as the root.

### BCR assembly and annotation

The IGHC gene type of each B cells was determined if the UMI count of that gene ≥ 3. For cells with more than one IGHC gene expressed, the UMI count of the major gene should be three times larger than the minor one. We assembled single-cell BCR contigs using TRUST4 (v1.0.6) [35], taking known IGHV/D/J/C sequences of Bactrian camels as the reference [13]. The raw FASTQ files were used as the input, and “--barcodeRange 0 15 + --umiRange 16 25 + --read1Range 40 -1” was set in compliance with the sequence composition of Read 1. Only barcodes in the 10x Genomics white list were preserved. We re-annotated the assembled contigs with IgBLAST (v1.10.0) [36], including the FR2 and CDR3 sequences in compliance with the IMGT standards [45]. Contigs with short alignment length (<150 bp), or non-productive sequences were removed. We determined VH/VHH contigs based on G49E/Q and L50R in complete FR2, which were the most discriminative sites [15]. For cells with more than one contig containing different FR2 or CDR3, the one with the highest coverage was chosen. If a cell contained both VH and VHH contigs, we assigned its IGHV type only when the coverage of the major type was three times larger than the minor type. BCR sequences and B cell transcriptome were matched based on the same cell barcodes.

### Camel immunization

We immunized two camels using BSA as an antigen for 4 times, with each immunization scheduled 14 days apart. Each immunization used an antigen dose of 0.5 mg (1 mg/ml), mixed with an equal volume of complete Freund’s adjuvant (initial immunization) or incomplete Freund’s adjuvant, and thoroughly emulsified for subcutaneous multi-point injection in the neck. Before immunization, 10 ml of whole blood was collected for serum titer determination using ELISA.

### ELISA

BSA was diluted in pH 9.6 carbonate-bicarbonate buffer to a concentration of 2 μg/ml, and 100 μl was added per well for coating overnight at 4°C. After discarding the coating solution, the plate was washed three times with 0.1% PBST (PBS with Tween 20), and then 300 μl of 5% skim milk (dissolved in 0.1% PBST) was added to each well for blocking at 37°C for 1 hour. After discarding the blocking solution, the plate was washed once with 0.1% PBST, and then 100 μl of different dilutions of antiserum (primary antibody, diluted in 5% skim milk) was added to each well at 37°C for 1 hour. Serum from unimmunized animals was used as the negative control, and 5% skim milk was used as the blank control. After washing the plate five times with 0.1% PBST, 100 μl HRP-conjugated goat anti-camel IgG (secondary antibody, NBbiolab, diluted 1:10,000) was added to each well and incubated at 37°C for 1 hour. After washing the plate five times with 0.1% PBST, 100 μl TMB substrate solution was added per well for color development and incubated at 37°C for 7 minutes. The reaction was stopped by adding 50 μl of stop solution (6M HCl), and the OD450 value was measured. A positive well was defined as a serum sample with an OD450 value greater than 3 times that of the unimmunized serum and a reading greater than 0.2. The antibody titer was defined as the maximum dilution of the positive wells.

### Differential expression analysis

We processed post-immunization scRNA-seq data with the same pipeline as we did for the pre-immunization data. Cell type annotations were transferred from the pre-immunization samples with Seurat’s reference transfer utility [25]. We performed differential expression analysis between pre- and post-immunization samples using the pseudobulk approach, which could account for the intrinsic variability of biological replicates and was shown superior to single-cell methods [40]. Pseudobulk UMI count matrices for major cell types and VHH+/VH+ populations were constructed with Libra (v1.0.0) [40], and DEGs were identified with DESeq2 (v1.30.1) [46]. The design formular for DESsq2 was ∼ *camel_ID* + *immunization_condition*, and the cutoff for DEGs was BH-adjusted *P* < 0.05. We used clusterProfiler (v3.16.0) [47] for functional over-representation analyses of DEGs, taking the Gene Ontology (GO) [48] of human orthologs as the database. Up- and down-regulated DEGs were analyzed separately, and the cutoff for significant GO terms was BH-adjusted *P* < 0.05.

### Data availability

The raw sequencing data generated in this study have been deposited in the Sequence Read Archive (SRA) database under accession code PRJNA997575 (https://www.ncbi.nlm.nih.gov/sra). The processed data have been deposited in the Gene Expression Omnibus (GEO) database under accession code GSE238082 (https://www.ncbi.nlm.nih.gov/geo). Source code used for analyzing the data are available at a public GitHub repository: https://github.com/zhenwang100/Abseq.

## Conflict of interest

The authors declare no competing interests.

## Acknowledgments

This work was supported by the National Natural Science Foundation of China (32070570), the Special Fund for Commercialization of Scientific and Research Findings in Inner Mongolia Autonomous Region (2021CG0021) and the National Key Research and Development Project (2020YFE0203300).

## Notes

### Competing Interest Statement

The authors have declared no competing interest.

