## Supplementary Materials for "Single-cell 5’ RNA sequencing of camelid peripheral B cells reveals cellular basis of heavy-chain antibody production"

Supplementary Table 1. Camel information.

| ID | Sex | Age | Location |
| --- | --- | --- | --- |
| C1 | Male | 5 years | Sonid Right Banner, Inner Mongolia |
| C2 | Male | 5 years | Sonid Right Banner, Inner Mongolia |
| C3 | Female | 4 years | Sonid Right Banner, Inner Mongolia |
| C4 | Female | 4 years | Sonid Right Banner, Inner Mongolia |

Supplementary Table 2. Summary of scRNA-seq data before immunization by CellRanger.

| **Sample ID** | C1 | C2 | C3 | C4 |
| --- | --- | --- | --- | --- |
| **Sequencing** | | | | |
| Number of Reads | 386,039,851 | 408,845,258 | 415,905,110 | 365,934,099 |
| Number of Short Reads Skipped | 0 | 0 | 0 | 0 |
| Valid Barcodes | 90.70% | 89.80% | 93.10% | 92.80% |
| Valid UMIs | 99.80% | 99.70% | 99.70% | 99.80% |
| Sequencing Saturation | 87.40% | 91.20% | 84.70% | 78.50% |
| Q30 Bases in Barcode | 97.20% | 97.30% | 96.90% | 96.90% |
| Q30 Bases in RNA Read | 91.40% | 91.20% | 93.30% | 93.40% |
| Q30 Bases in RNA Read 2 | 93.30% | 92.90% | 92.00% | 91.50% |
| Q30 Bases in UMI | 96.60% | 96.70% | 96.20% | 96.10% |
| **Mapping** | | | | |
| Reads Mapped to Genome | 84.70% | 82.70% | 87.20% | 87.50% |
| Reads Mapped Confidently to Genome | 83.00% | 81.10% | 85.40% | 85.80% |
| Reads Mapped Confidently to Intergenic Regions | 5.50% | 5.50% | 5.90% | 5.40% |
| Reads Mapped Confidently to Intronic Regions | 6.80% | 6.80% | 6.60% | 7.70% |
| Reads Mapped Confidently to Exonic Regions | 70.80% | 68.80% | 72.90% | 72.70% |
| Reads Mapped Confidently to Transcriptome | 65.00% | 63.30% | 68.20% | 67.30% |
| Reads Mapped Antisense to Gene | 3.20% | 3.00% | 2.10% | 2.60% |
| **Cells** | | | | |
| Estimated Number of Cells | 6,834 | 5,725 | 7,060 | 10,136 |
| Fraction Reads in Cells | 93.50% | 95.10% | 90.60% | 90.90% |
| Mean Reads per Cell | 56,488 | 71,414 | 58,910 | 36,102 |
| Median UMI Counts per Cell | 2,737 | 2,761 | 4,034 | 2,984 |
| Median Genes per Cell | 1,274 | 1,274 | 1,725 | 1,448 |
| Total Genes Detected | 15,156 | 14,978 | 15,422 | 15,593 |

Supplementary Table 3. Summary of scRNA-seq data after immunization by CellRanger.

| **Sample ID** | C3-42 day | C3-56 day | C4-42day | C4-56 day |
| --- | --- | --- | --- | --- |
| **Sequencing** | | | | |
| Number of Reads | 414,579,447 | 480,443,763 | 411,728,306 | 458,571,083 |
| Number of Short Reads Skipped | 0 | 0 | 0 | 0 |
| Valid Barcodes | 93.10% | 93.00% | 92.60% | 93.00% |
| Valid UMIs | 99.40% | 99.60% | 99.80% | 99.60% |
| Sequencing Saturation | 82.00% | 90.00% | 78.60% | 88.20% |
| Q30 Bases in Barcode | 97.00% | 96.90% | 97.00% | 96.80% |
| Q30 Bases in RNA Read | 92.80% | 92.90% | 93.30% | 92.90% |
| Q30 Bases in RNA Read 2 | 89.30% | 89.80% | 92.00% | 91.10% |
| Q30 Bases in UMI | 96.20% | 95.90% | 96.20% | 95.90% |
| **Mapping** | | | | |
| Reads Mapped to Genome | 83.00% | 86.10% | 85.00% | 85.30% |
| Reads Mapped Confidently to Genome | 81.50% | 84.60% | 83.20% | 83.80% |
| Reads Mapped Confidently to Intergenic Regions | 5.40% | 5.00% | 5.30% | 5.30% |
| Reads Mapped Confidently to Intronic Regions | 7.80% | 6.70% | 8.00% | 6.70% |
| Reads Mapped Confidently to Exonic Regions | 68.40% | 73.00% | 69.90% | 71.70% |
| Reads Mapped Confidently to Transcriptome | 63.60% | 68.00% | 65.20% | 67.30% |
| Reads Mapped Antisense to Gene | 2.00% | 2.30% | 2.10% | 2.00% |
| **Cells** | | | | |
| Estimated Number of Cells | 8,776 | 5,784 | 9,784 | 7,395 |
| Fraction Reads in Cells | 90.50% | 96.50% | 94.70% | 93.00% |
| Mean Reads per Cell | 47,240 | 83,064 | 42,082 | 62,011 |
| Median UMI Counts per Cell | 3,370 | 3,818 | 3,590 | 3,335 |
| Median Genes per Cell | 1,518 | 1,635 | 1,569 | 1,464 |
| Total Genes Detected | 15,556 | 15,234 | 15,638 | 15,310 |

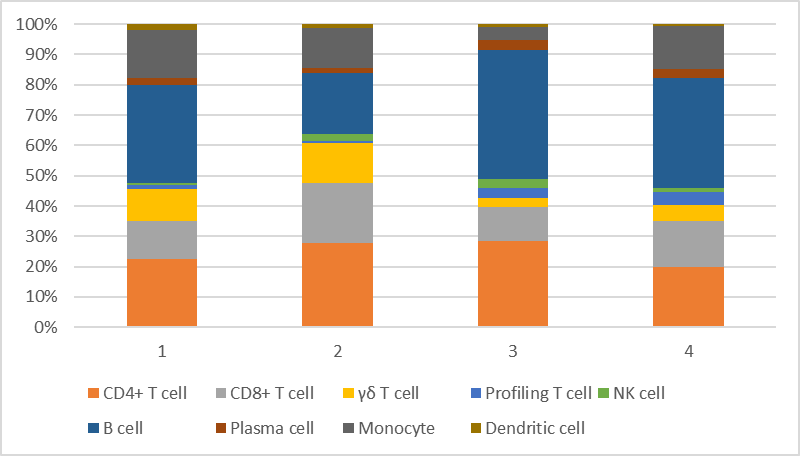

Supplementary Figure 1. PBMC compositions in the four camels.

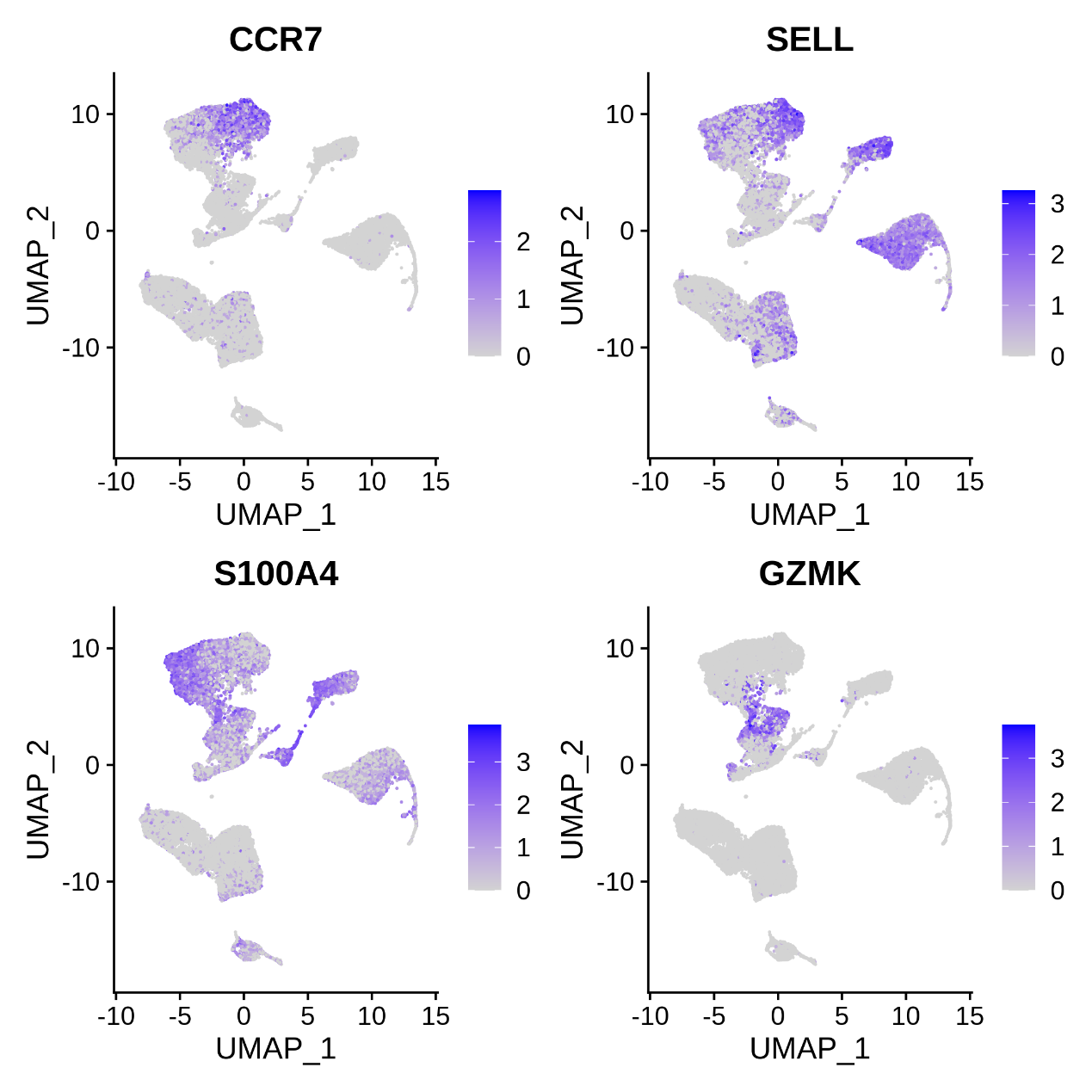

Supplementary Figure 2. Gene markers for T cell subsets. CCR7 and SELL are markers for naive T cells, while S100A4 and GZMK are markers for memory T cells.

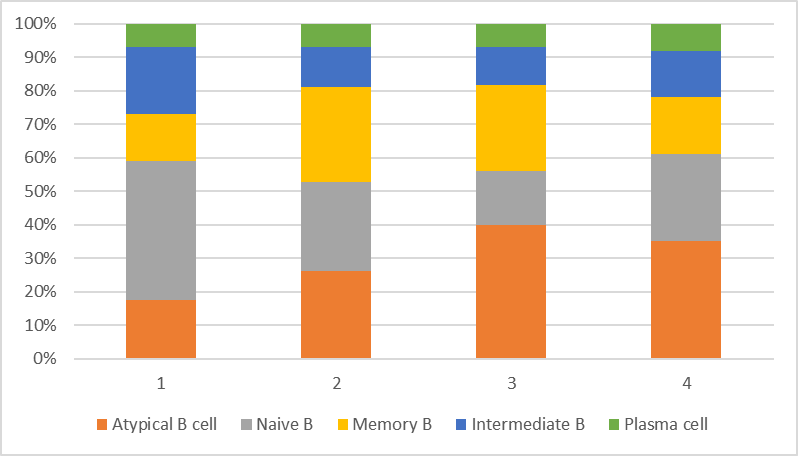

Supplementary Figure 3. B cell compositions in the four camels.

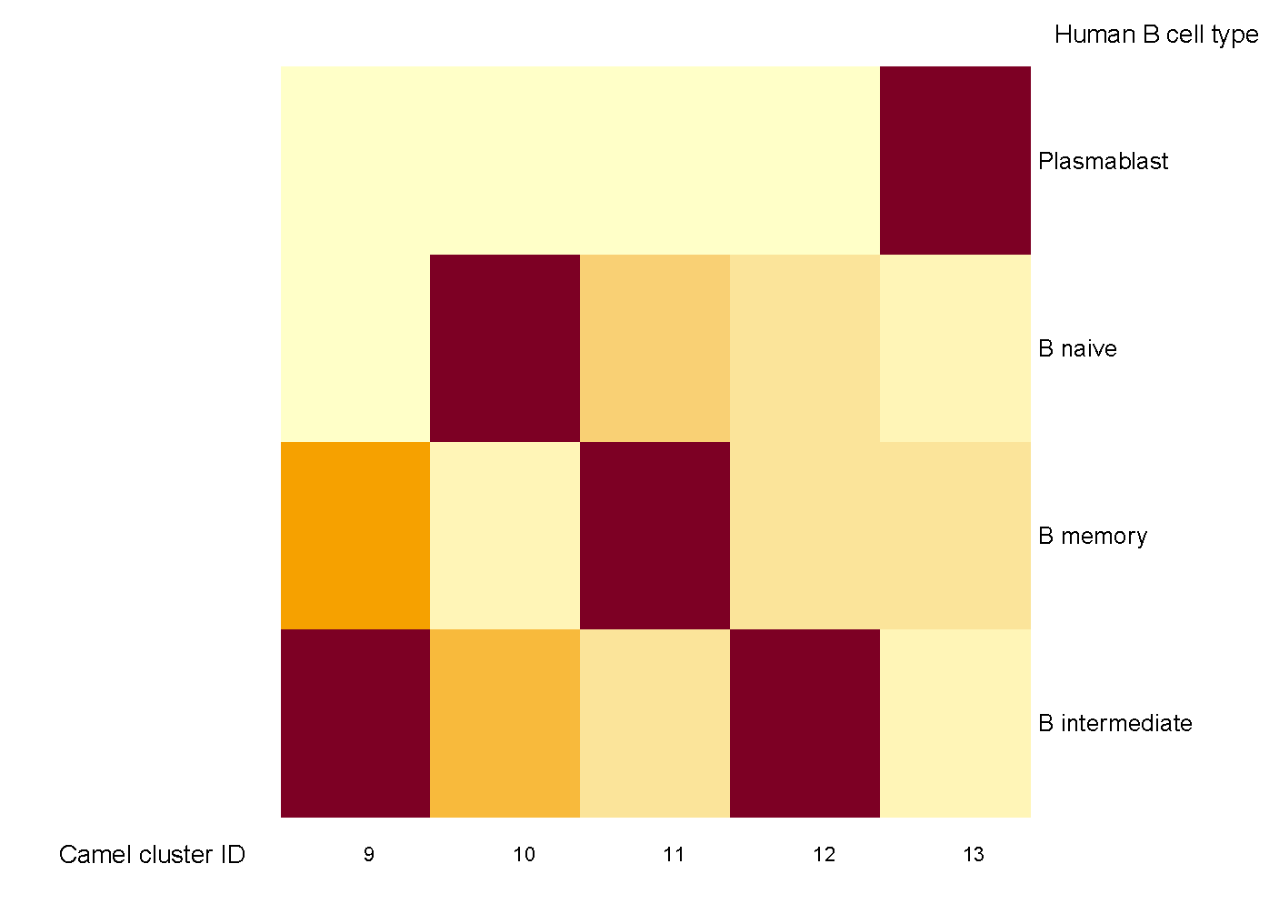

Supplementary Figure 4. Mapping camel B cells to Seurat’s human PBMC reference. Cluster 10, 11, 12 and 13 correspond to human naive B cells, memory B cells, intermediate B cells and plasmablasts, respectively. Note Cluster 9 (atypical B cells) lacks corresponding annotation in human PBMCs.

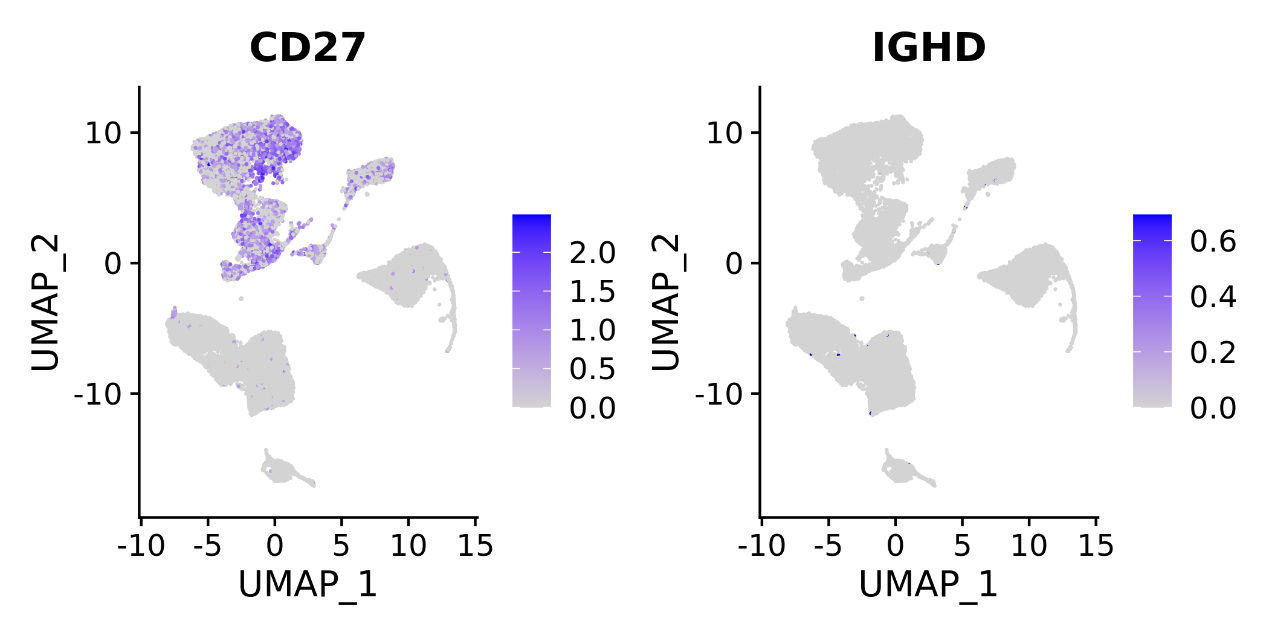

Supplementary Figure 5. Expression of CD27 and IGHD. CD27 is a marker gene for human memory B cells, and IGHD is a marker gene for human naive B cells. Both genes lack expression in all camel B cells.

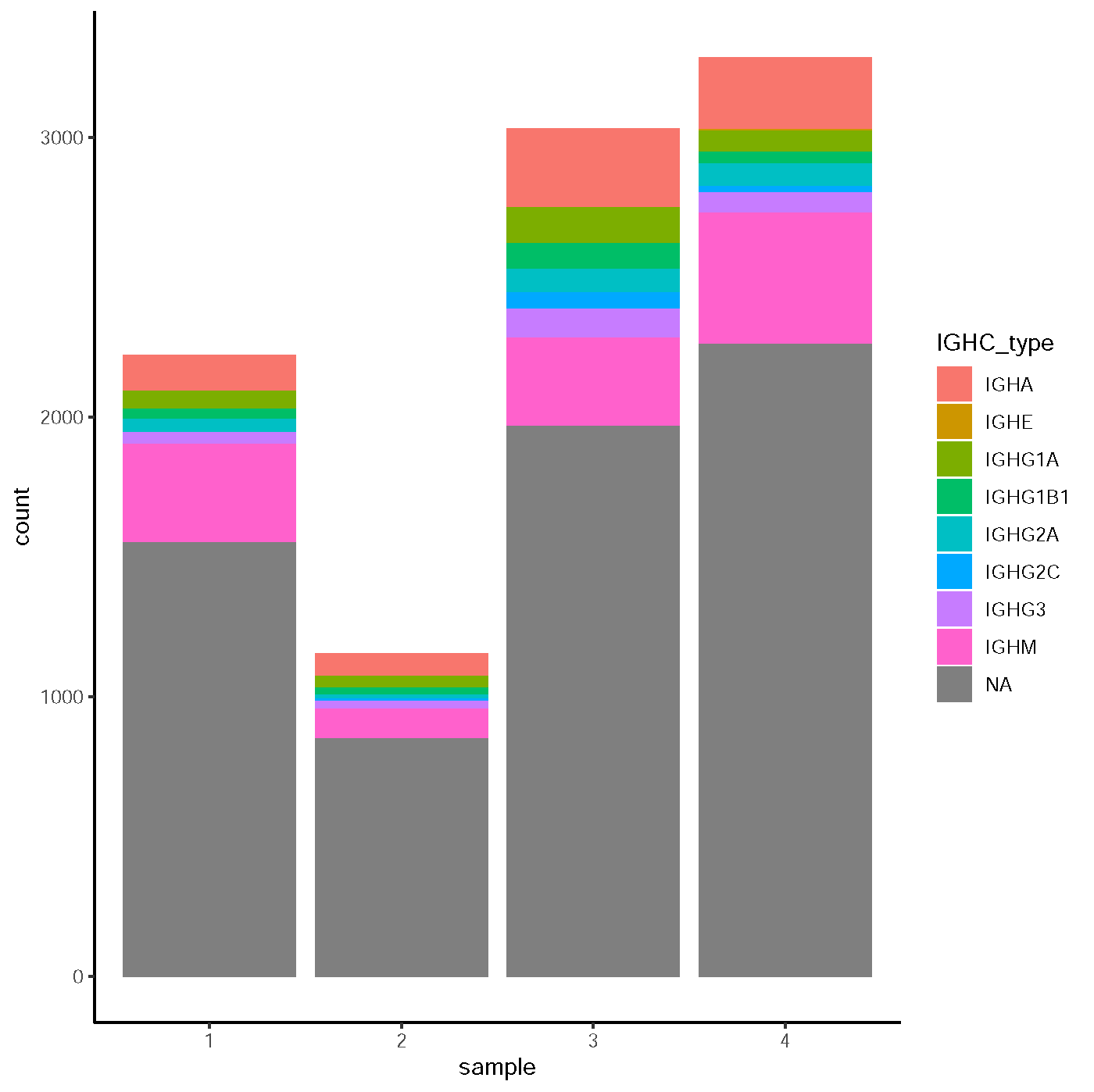

Supplementary Figure 6. Number of B cells with different IGHC types in the four camels.

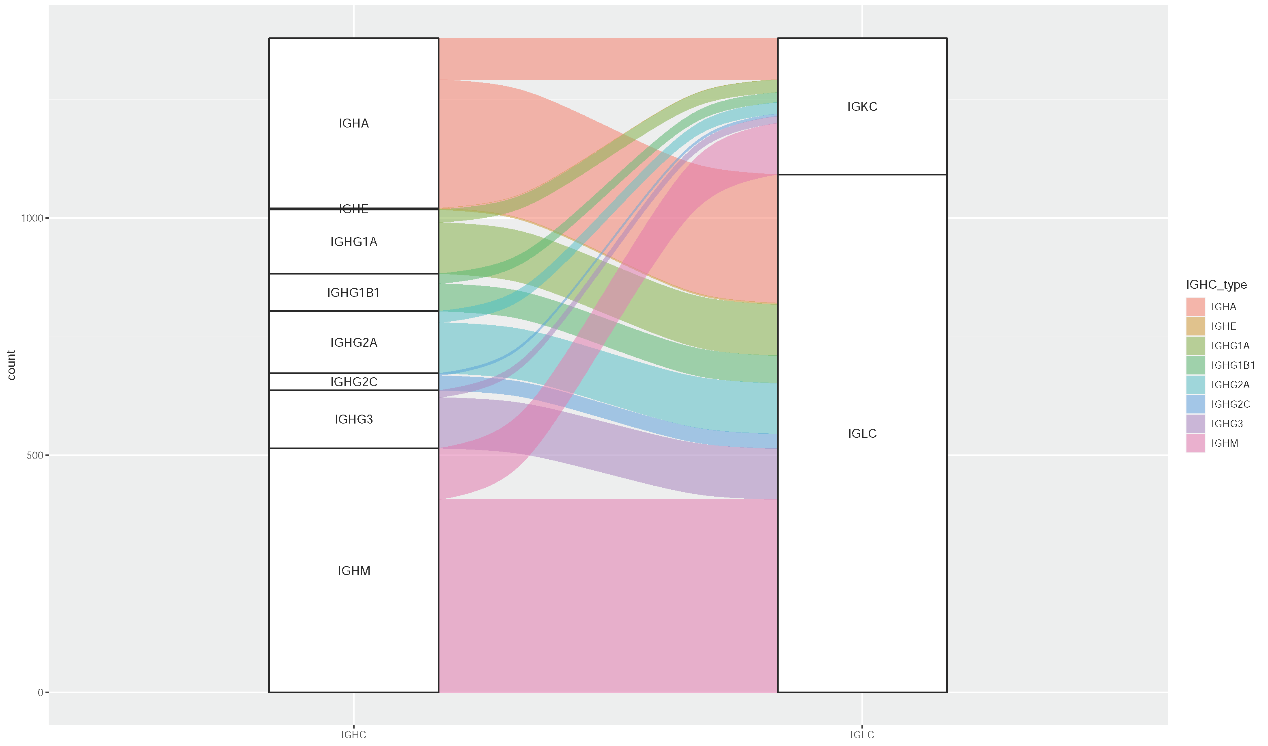

Supplementary Figure 7. Association between IGHC and IGLC/IGKC. In B cells express HCAb genes (IGHG2/3), there are also expression of light chain genes (IGLC/IGKC).

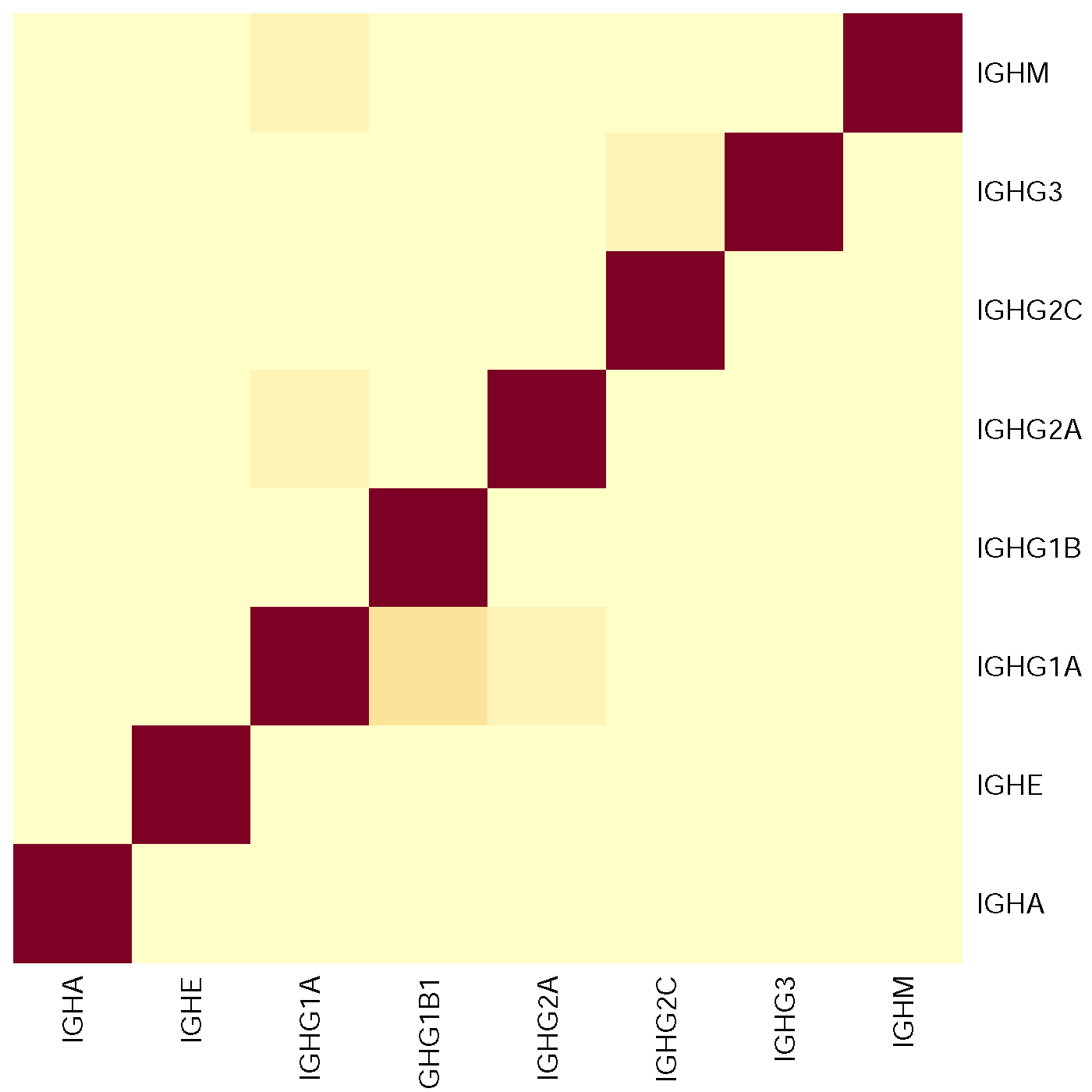

Supplementary Figure 8. Comparison of IGHC types recovered by gene expression and TRUST4 assembly.

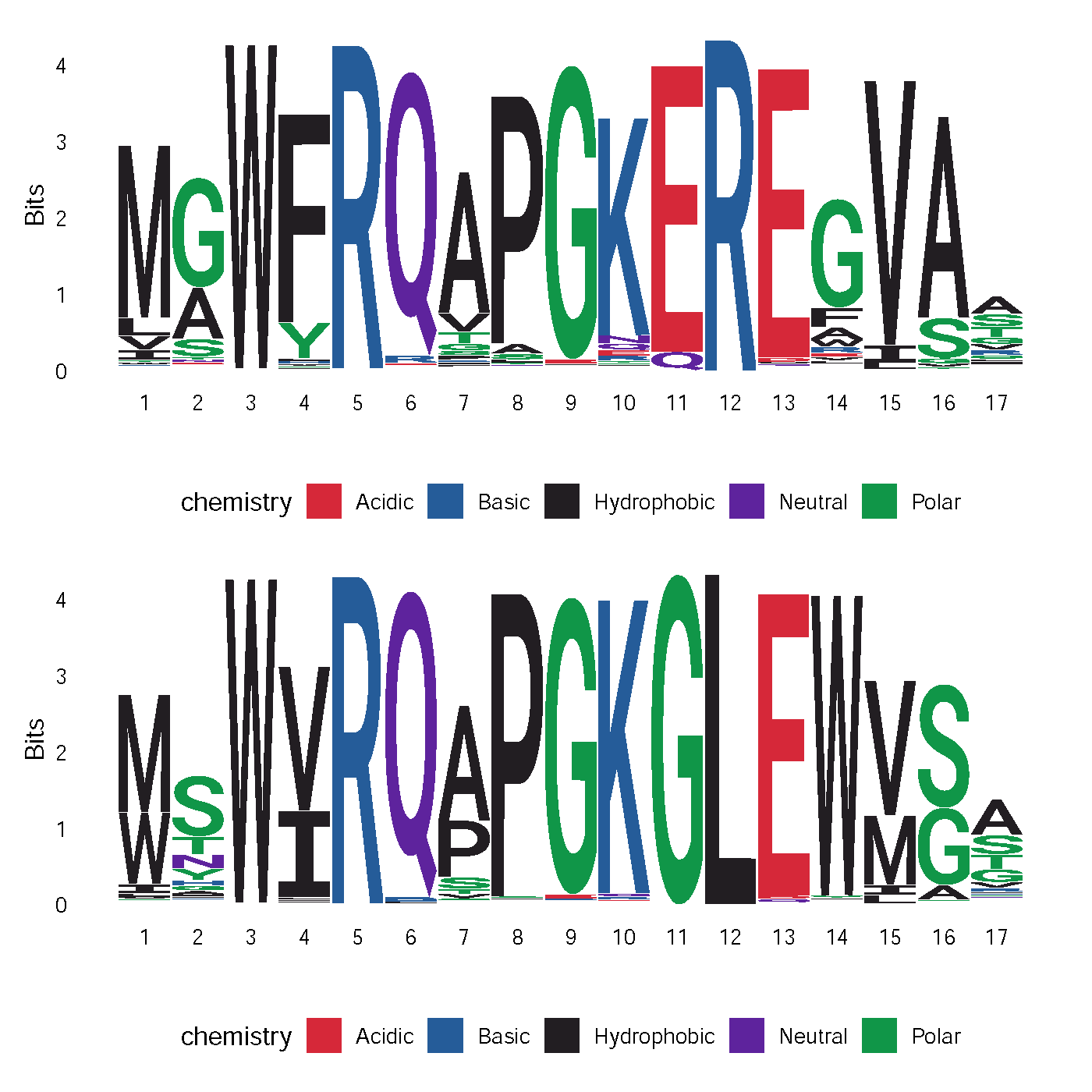

Supplementary Figure 9. Sequence logo of FR2 in VHH (upper) and VH (lower). The most discriminative sites are G49E/Q and L50R (Site 11 and 12 in the figure).

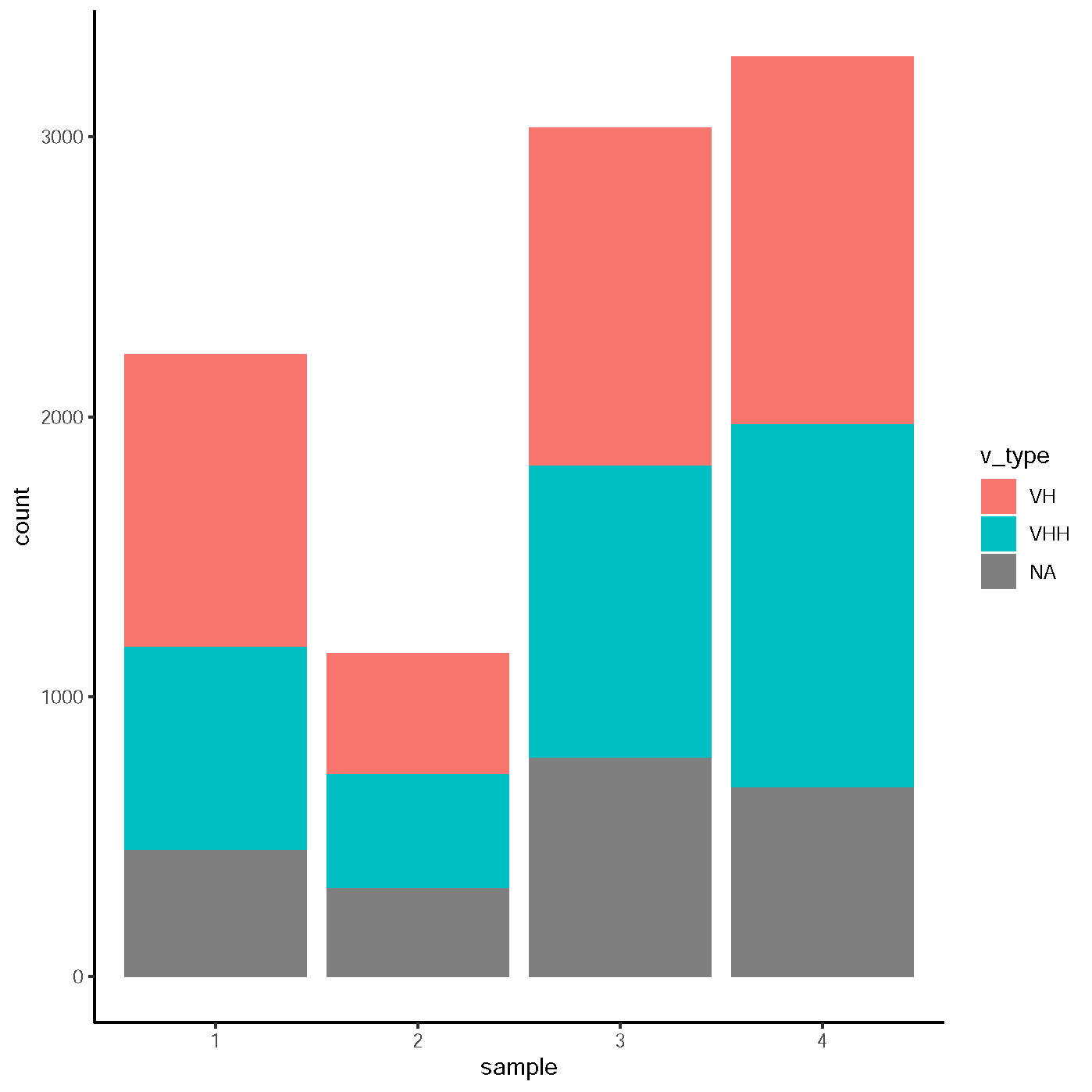

Supplementary Figure 10. Number of B cells with different IGHV types in the four camels.

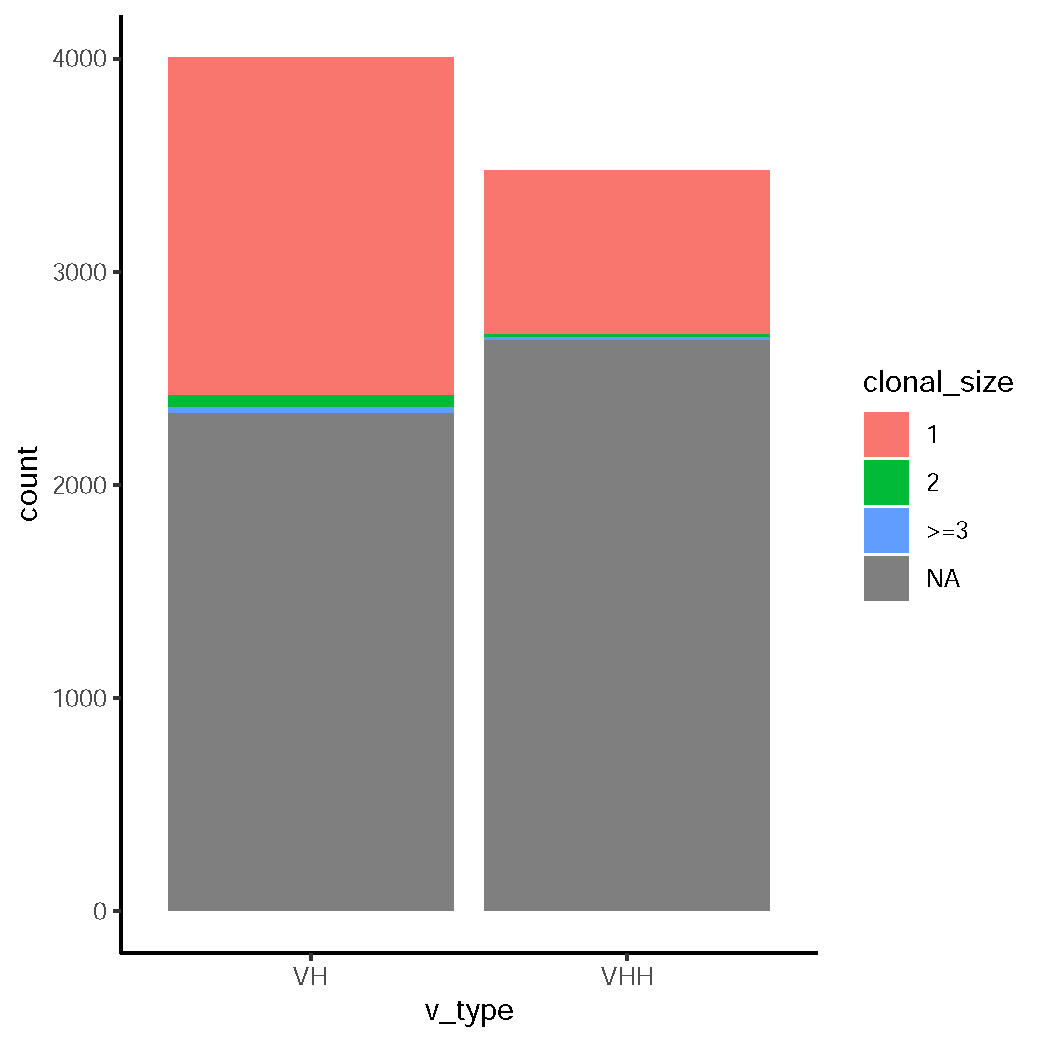

Supplementary Figure 11. Clonal size in VH+ and VHH+ B cells. The clonotypes are determined by CDR3 sequences.

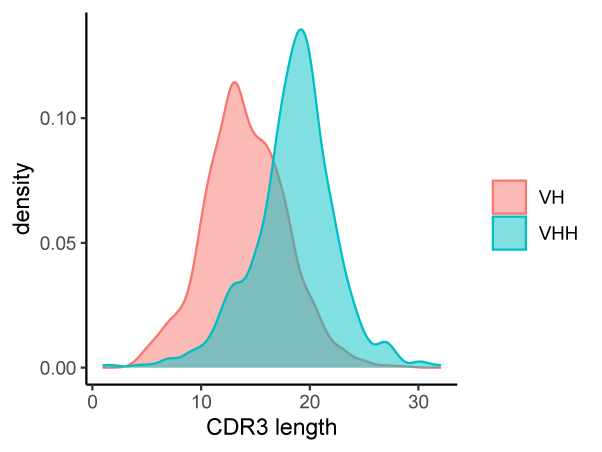

Supplementary Figure 12. Comparison of CDR3 length (amino acids) between VH and VHH. The CDR3 length of VHH was significantly longer than that of VH.

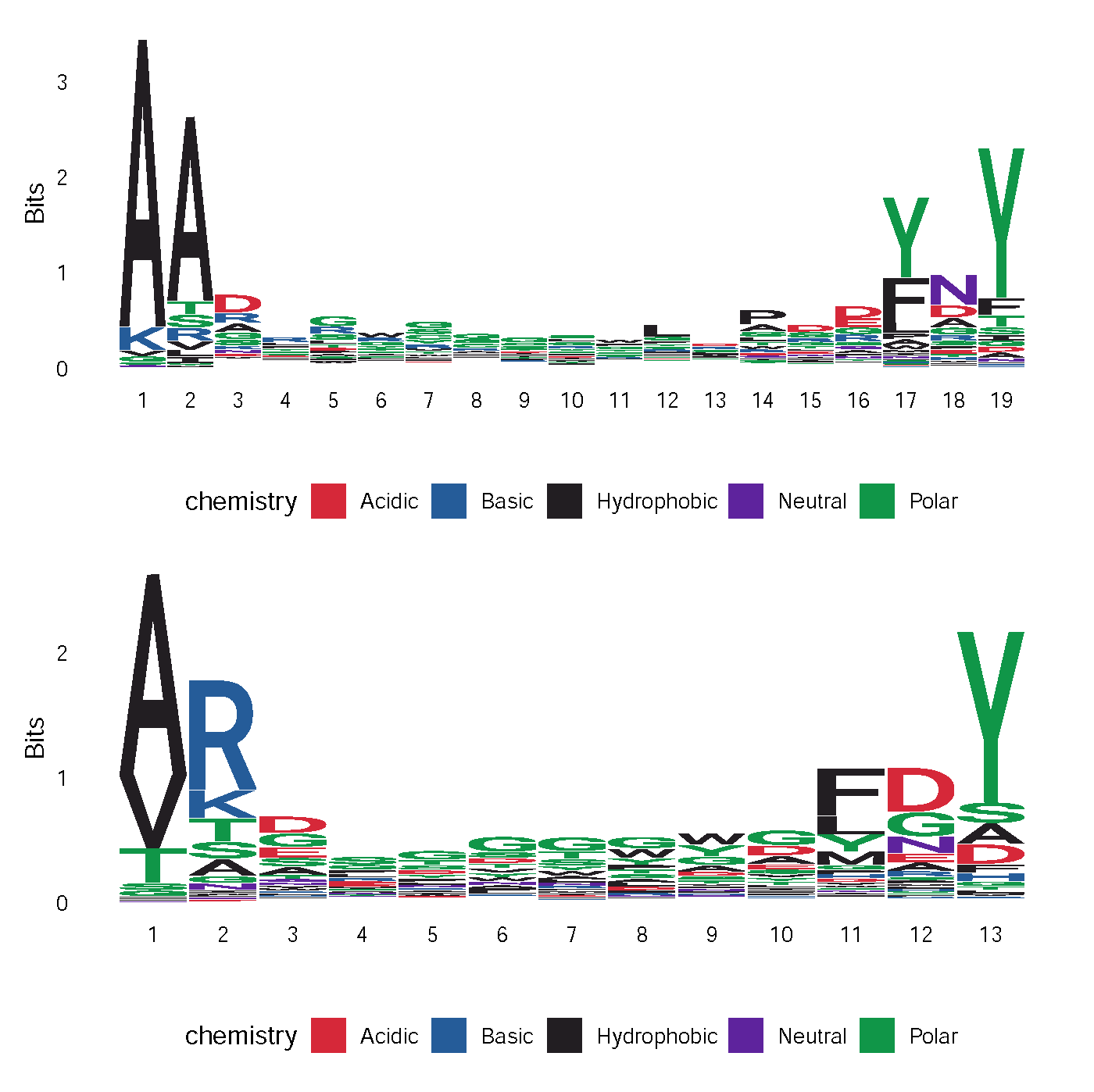

Supplementary Figure 13. Sequence logo of CDR3 in VHH (upper) and VH (lower). The most common length for VHH (19) and VH (13) are plotted.

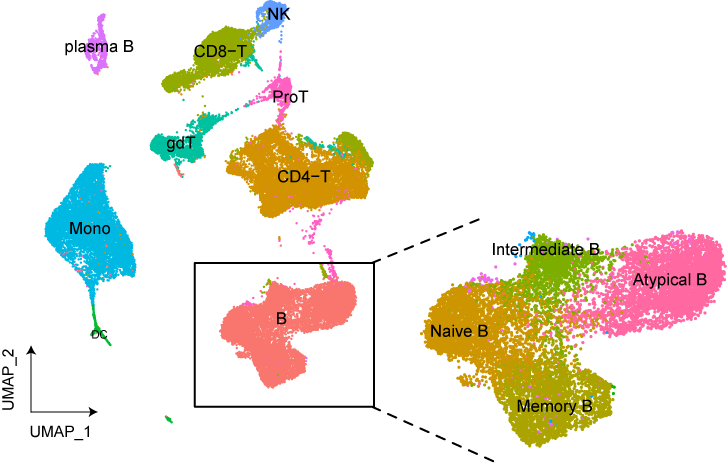

Supplementary Figure 14. UMAP visualization of PBMC clustering after immunization. The cell type annotations are transferred from PBMCs before immunization. The annotations of B cell subtypes are indicated in the inserted panel.

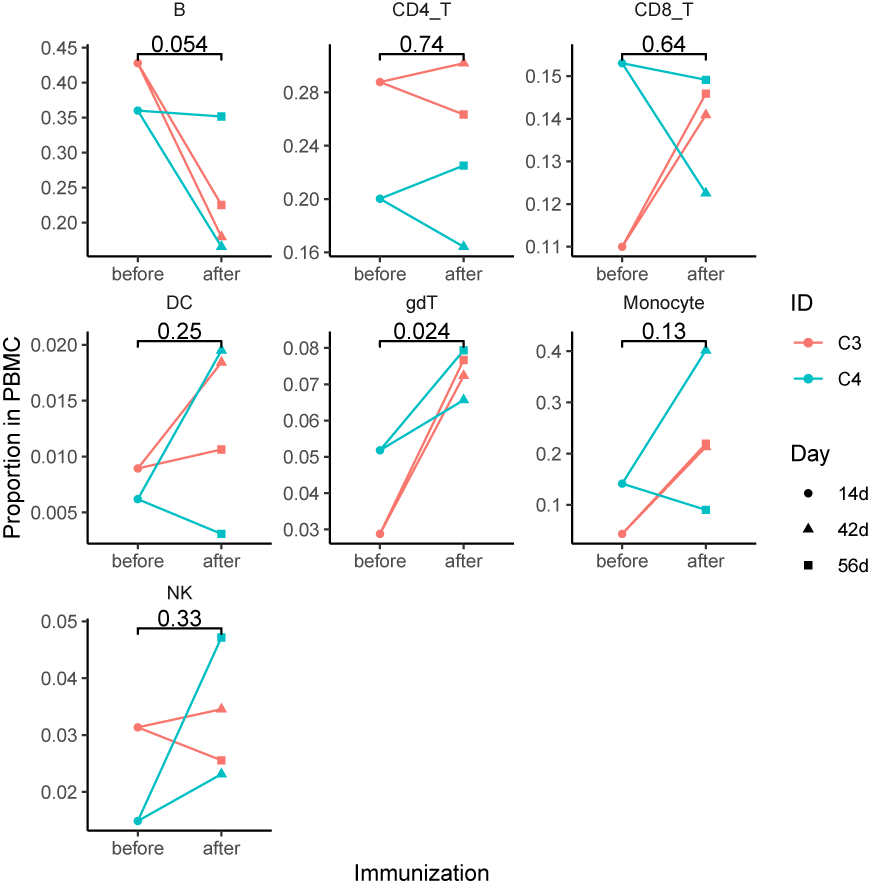

Supplementary Figure 15. Proportion of major cell types in PBMCs before and after immunization. *P*-values are calculated for comparisons (two-sided *t*-test).

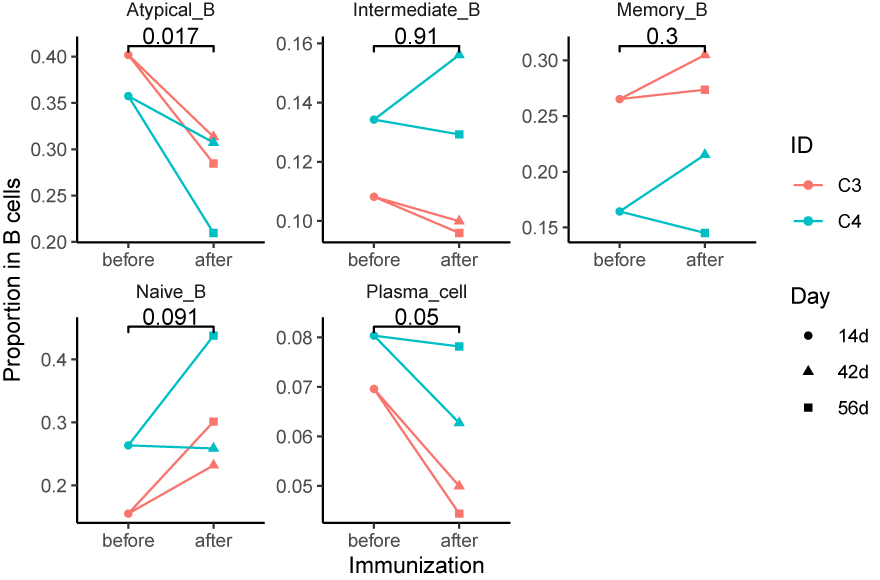

Supplementary Figure 16. Proportion of B subtypes in all B cells before and after immunization. *P*-values are calculated for comparisons (two-sided *t*-test).

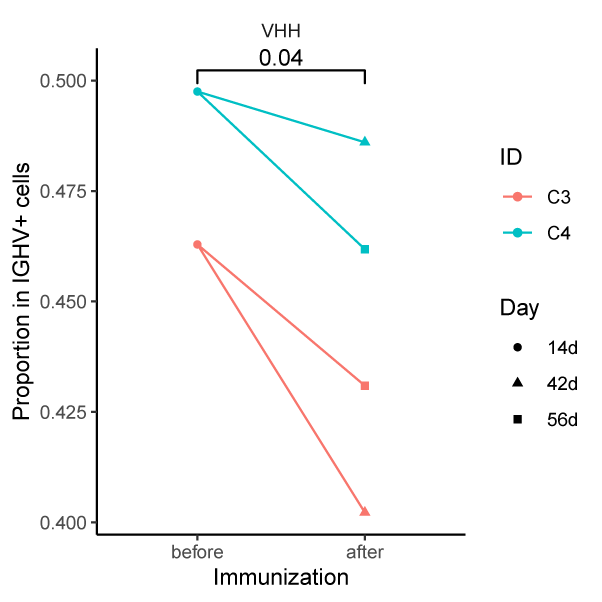

Supplementary Figure 17. Proportion of VHH+ cells in all IGHV+ cells before and after immunization. *P*-values are calculated for comparisons (two-sided *t*-test).

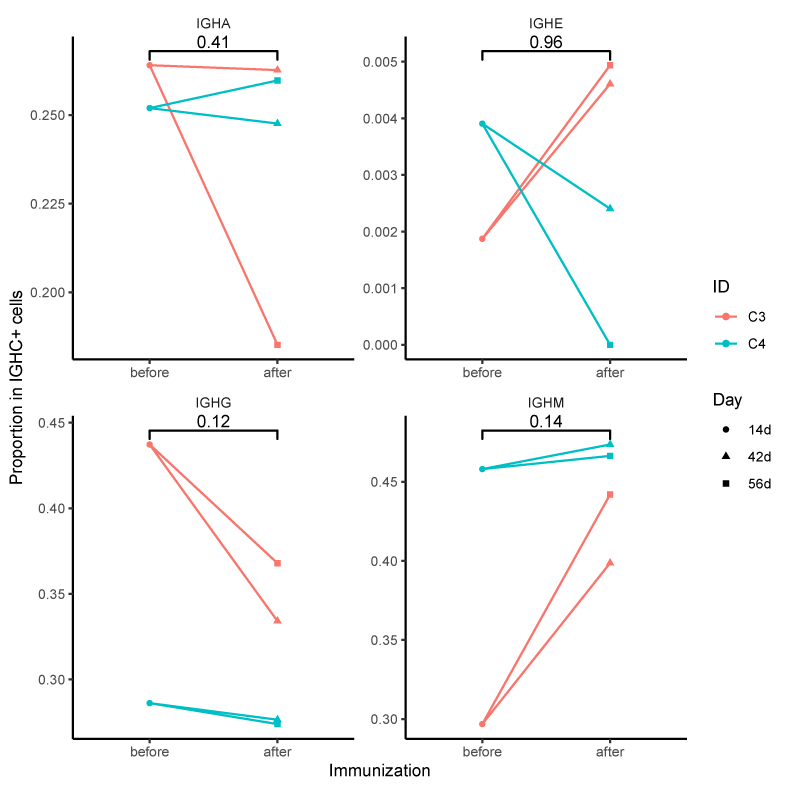

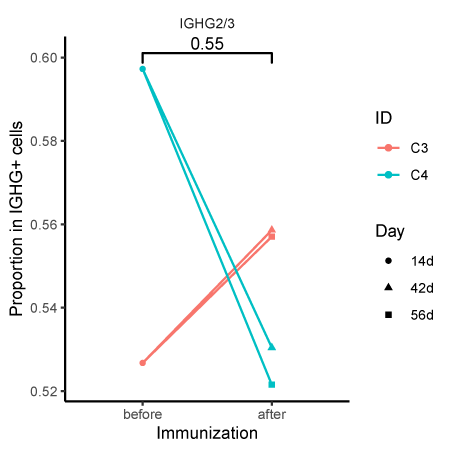

Supplementary Figure 18. Proportion of IGHC types before and after immunization. (Upper) Major types in all IGHC+ cells. (Lower) IGHG2/3 types in all IGHG+ cells. *P*-values are calculated for comparisons (two-sided *t*-test).

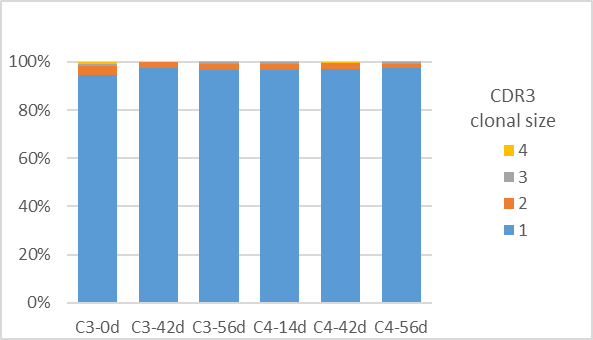

Supplementary Figure 19. Proportion of B cells with different clonal size during immunization. The clonotypes are determined by IGHV CDR3 sequences.
